# Electrochemical Deformation of PEDOT:PSS Drives Mechanosensitive Cell Activation

**DOI:** 10.64898/2026.08.12.744395

**Authors:** Anna Fiene Müller, Franziska Wasner, Ryan W. Crisp, Julien Bachmann, Vicente Duran-Toro, Danijela Gregurec

**Affiliations:** Biointerfaces Lab, Department of Chemistry and Pharmacy, Friedrich-Alexander Universität Erlangen- Nürnberg, Henkestrasse 91, 91052 Erlangen, Germany; Department of Chemistry and Pharmacy, Section of Materials Chemistry, Chair of Chemistry of Thin Film Materials, Friedrich-Alexander-Universität Erlangen-Nürnberg, Cauerstr. 3, 91058 Erlangen, Germany

**Keywords:** PEDOT:PSS, conducting polymers, electrochemical actuation, mechanotransduction, mechanosensitive ion channels, bioelectronic interfaces, calcium signaling

## Abstract

Conducting polymers are widely used in bioelectronic interfaces because of their mixed ionic-electronic conductivity, mechanical compliance, and compatibility with biological systems. However, their electrochemically driven structural dynamics have received little attention as a mechanism for mechanical cell stimulation. Here, we show that electrochemical actuation of poly(3,4-ethylenedioxythiophene):polystyrenesulfonate (PEDOT:PSS) generates mechanical cues capable of activating endogenous mechanosensitive pathways in HEK293T cells. Transparent PEDOT:PSS films deposited on ITO exhibited a heterogeneous granular morphology and underwent potential-dependent microscopic deformation during electrochemical modulation. Direct optical tracking revealed displacement of the polymer boundary, with structural changes occurring preferentially in polymer-dense regions and propagating toward the film edge. When HEK293T cells were cultured directly on PEDOT:PSS, repeated electrochemical stimulation at -240 mV produced reproducible intracellular Ca^2+^ responses. Pharmacological inhibition with GsMTx4 attenuated the calcium response, whereas blockade of voltage-gated sodium channels with tetrodotoxin largely preserved it, supporting the involvement of mechanosensitive pathways in the cellular response. These findings identify PEDOT:PSS as an electromechanical biointerface in which electrochemical modulation can introduce a mechanical component alongside the established electrical function of the interface. This mechanical contribution should therefore be considered when interpreting cellular responses to conducting polymer- based electrical stimulation and provides a basis for engineering bioelectronic interfaces that deliberately couple electrical control with mechanotransduction.

## INTRODUCTION

Conducting polymers have become indispensable materials for bioelectronic interfaces because they combine efficient electronic transport with ionic conductivity and mechanical properties compatible with biological tissues. Among them, poly(3,4-ethylenedioxythiophene):polystyrenesulfonate (PEDOT:PSS) has emerged as one of the most widely used materials for neural interfaces,(1) and electrophysiological devices owing it to its high electrical conductivity,(2) mechanical properties,(3) and compatibility with microfabrication.(4) Despite these advances, PEDOT:PSS has been predominantly exploited as an electrical biointerface, whereas its electrochemically driven structural changes have received little attention as a potential mechanism for mechanical stimulation of living cells.

During electrochemical reduction, PEDOT:PSS undergoes ion exchange in which hydrated cations enter the polymer network as PEDOT is reduced, maintaining charge compensation with the negatively charged PSS phase. This process produces reversible swelling and contraction of the polymer matrix that has been extensively investigated in the context of electrochemical energy storage, soft actuators, and electrochromic devices.(5) Despite the well-established coupling between electrochemical state and mechanical deformation, it remains unknown whether these dynamic structural changes can generate sufficient mechanical strain at the cell-material interface to activate mechanosensitive signaling pathways.

Mechanosensitive ion channels convert mechanical forces into intracellular biochemical signals and play central roles in processes including touch sensation, hearing, vascular regulation, and cellular homeostasis.(6, 7) Mechanical deformation of the plasma membrane can alter membrane tension and activate mechanosensitive channels, including PIEZO1 and TRPV4, resulting in intracellular Ca^2+^ signaling. Because HEK293T cells endogenously express mechanosensitive ion channels while exhibiting limited voltage-gated excitability,(8-10) they provide a suitable model for distinguishing mechanically induced calcium signaling from conventional electrical excitation.

To investigate whether electrochemically induced deformation of PEDOT:PSS, such as swelling under negative reducing potentials(5, 11), can mechanically activate cells, we developed a transparent PEDOT:PSS electrode compatible with simultaneous electrochemical modulation and live cell calcium imaging. Integration of this platform with pharmacological inhibition of mechanosensitive and voltage gated sodium channels enabled us to distinguish their contributions to the cellular response. We show that electrochemical modulation produces spatially heterogeneous deformation of PEDOT:PSS that is accompanied by activation of endogenous mechanosensitive pathways in adherent HEK293T cells. These findings establish PEDOT:PSS as an electromechanical biointerface and reveal that electrochemical stimulation through a conducting polymer can introduce a mechanical component at the cell-material interface alongside its established electrical function.

## MATERIALS AND METHODS

### Materials

For electrode fabrication 2-propanol 98% (Carl Roth), 3-glycidoxypropyltrimethoxysilane (GOPS) 98% (Sigma-Aldrich), 4 -dodecylbenzenesulfonic acid (DBSA) 95% (Sigma-Aldrich), acetone 98% (Sigma-Aldrich), ethanol absolute (Fischer Scientific), ethylene glycol 99% (Sigma-Aldrich), indium tin oxide coated glass slide (ITO glass, 70-100 Ω/sq, Sigma-Aldrich), poly(3,4-ethylenedioxythiophene) polystyrene sulfonate 1.3 % (PEDOT:PSS; Ossila) and Toluene 99% (Carl Roth) were purchased and used as delivered. For electrochemical characterization Potassium chloride (KCl 99%, Sigma-Aldrich) was used as received.

For cell cultures Dulbecco’s modified Eagle’s Medium (DMEM), fetal bovine serum 10 % (FBS), phosphate-buffered saline (PBS, pH 7.4), Trypan Blue (0.4%), TrypLE Express, and penicillin-streptomycin were purchased from Fisher Scientific. For calcium imaging experiments, BioTrack 609 Ca^2+^ AM (Sigma Aldrich), Fluo-4 AM (Fisher Scientific), GsMTx4 trifluoroacetate, tetrodotoxin (TTX, Biomol GmBH)), and Tyrode solution (Thermo Fisher) were used. Ultrapure Milli-Q water (18.2 MΩ cm) was used throughout the study.

### ITO/PEDOT:PSS electrodes fabrication

ITO-coated glass was cut into 25 mm × 8 mm pieces and sequentially cleaned by sonication in acetone, 2-propanol, and deionized (DI) water for 10 min each, following the protocol described by Osazuwa et al.(12) The substrates were dried under compressed air and exposed to UV light for 30 min, as previously described.(13) For surface functionalization, a solution containing 1% (v/v) (3-glycidyloxypropyl)trimethoxysilane (GOPS) in anhydrous toluene was prepared. The ITO substrates were immersed in the GOPS solution and kept in a desiccator overnight. The following day, the substrates were rinsed with toluene, dried under compressed air, and stored in a desiccator until further use.

The PEDOT:PSS coating solution was prepared by combining the aqueous PEDOT:PSS dispersion with ethylene glycol (5%, v/v) and dodecylbenzenesulfonic acid (DBSA; 0.25%, w/v), as previously described.(13) For dip coating, GOPS-functionalized ITO substrates were immersed in the PEDOT:PSS solution and immediately withdrawn. One to five PEDOT:PSS layers were deposited by repeating the dip-coating procedure, with annealing at 110 °C for 1 min between successive coatings. Following deposition, the coated substrates were stored in a desiccator overnight. The PEDOT:PSS-coated substrates were subsequently immersed in absolute ethanol for 3 h, followed by immersion in Milli-Q water overnight. The substrates were then dried and stored at 4 °C until use.

### Electrode characterization

Atomic force microscopy (AFM; Cypher S, Asylum Research, Oxford Instruments) was performed at room temperature in air-tapping mode using a Track60 probe (spring constant, 1.6 N m^-1^) to characterize the surface topography of the PEDOT:PSS films. Fourier transform infrared spectroscopy (FTIR; ALPHA, Bruker) was performed in attenuated total reflection (ATR) mode over the range 400 to 4000 cm^-1^.

Cyclic voltammetry (CV) and electrochemical impedance spectroscopy (EIS) were performed using a three-electrode electrochemical cell connected to an EmStat4X potentiostat (PalmSens BV). PEDOT:PSS-coated ITO served as the working electrode, with Ag/AgCl and Pt wire used as the reference and counter electrodes, respectively. Measurements were performed in 10 mM KCl. For CV, five scans were recorded between -0.8 and +0.8 V at a scan rate of 5 mV s^-1^ and a potential step of 0.05 V. EIS measurements were performed over a frequency range of 0.1 Hz-100 kHz using a potential range of 0-0.01 V, as previously described.(12)

For measurements in the presence of cells, PEDOT:PSS-coated ITO electrodes were autoclaved at 121 °C for 15 min before cell seeding as described below. Open-circuit potentiometry (OCP) was first recorded for 2 min with a sampling interval of 0.1 s to determine the resting potential of electrodes containing HEK293T cells at approximately 70% confluency. Subsequently, multistep amperometry was performed after a 30 s equilibration period. Three consecutive stimulation cycles were applied, each consisting of three 30 s potential levels. The first and third levels were maintained at the OCP determined immediately before stimulation, whereas the second level was set to -240 mV. This stimulation sequence was subsequently used for Ca^2+^ imaging experiments to provide a reproducible potential protocol across cell-seeded electrodes.

### HEK293T cells cultures on ITO/PEDOT:PSS electrodes

Human embryonic kidney HEK293T cells (ATCC, CRL-3216) were maintained in Dulbecco’s modified Eagle’s medium (DMEM) supplemented with 10% fetal bovine serum (FBS) and 1% penicillin-streptomycin. Cells were cultured at 37 °C in a humidified atmosphere containing 5% CO_2_ and routinely passaged using TrypLE Express upon reaching confluence. Low-passage cells were used throughout the experiments.

Three-layer PEDOT:PSS-coated ITO working electrodes were sterilized and placed in 6-well plates. HEK293T cells were seeded directly onto the PEDOT:PSS surface at a density of 1.5 × 10^5^ cells cm^-2^ in 3 mL of complete DMEM and incubated for 24 h at 37 °C and 5% CO_2_.

For Ca^2+^ imaging, cells were loaded with either Fluo-4 AM or BioTracker 609. Both fluorescent Ca^2+^ indicators have dissociation constants in the submicromolar range (K_d_ = 0.345 μM for Fluo-4 AM and 0.19 μM for BioTracker 609). The indicators were diluted to a final concentration of 5 μM in prewarmed 1× Tyrode’s solution. Culture medium was replaced with the indicator-containing Tyrode’s solution, and cells were incubated for 30 min at 37 °C and 5% CO_2_. Cells were subsequently washed carefully with prewarmed Tyrode’s solution to remove excess indicator while minimizing cell detachment.

For pharmacological inhibition experiments, Ca^2+^ indicator-loaded HEK293T cells were incubated with 1 µM GsMTx4 or 1 µM TTX in Tyrode’s solution for 5 min at 37 °C and 5% CO_2_, following our previously established protocol.(14, 15)

### Calcium imaging

The three-electrode electrochemical setup described above was used for Ca^2+^ imaging, with Tyrode’s solution containing 1.8 mM Ca^2+^ used as the electrolyte instead of KCl. The cell-seeded PEDOT:PSS/ITO working electrode was inverted and inserted into the horizontal electrochemical cell (Figure S2a), with the PEDOT:PSS surface facing the imaging objective. The chamber was filled with Tyrode’s solution to completely cover the electrode surface, resulting in a final working volume of 5 mL.

Live-cell fluorescence imaging was performed using an inverted microscope (IX73, Olympus) equipped with an ORCA-Flash4.0 V3 camera (Hamamatsu Photonics). BioTracker 609- and Fluo-4 AM-loaded cells were imaged using the corresponding fluorescence channels. Images were acquired at 4 frames s^-1^ with an exposure time of ≤200 ms. Illumination intensity was maintained at ≤15% to minimize photobleaching and pixel saturation. During imaging, the three-cycle potential protocol described above was applied, with each cycle consisting of 30 s at the experimentally determined OCP, 30 s at -240 mV, and 30 s at OCP. Recorded image sequences were analyzed in ImageJ by manually defining cellular regions of interest (ROIs). Fluorescence traces were subsequently processed in MATLAB using the moving-average filtering and ΔF/F_0_ analysis procedure previously described,(14) with parameters adapted to the present acquisition rate. For responder analysis, ΔF/F_0_ traces were binned over four consecutive frames, and cells were classified as responsive when the fluorescence signal exceeded five times the standard deviation of the prestimulation baseline (5σ). Fifty cells were randomly selected from each of six independently prepared cell-seeded electrodes per condition using a fixed randomization seed, resulting in 300 cells analyzed per condition.

## RESULTS AND DISCUSSION

### Engineering PEDOT:PSS biointerface for electromechanical cell stimulation

To investigate whether electrochemically driven deformation of PEDOT:PSS can mechanically stimulate living cells, we first established a transparent PEDOT:PSS biointerface compatible with both electrochemical modulation and fluorescence imaging. PEDOT:PSS films were deposited onto indium tin oxide (ITO) glass substrates by dip-coating and characterised using atomic force microscopy (AFM) and Fourier-transform infrared spectroscopy (FTIR) (Figure 1a,b, Figure S1).

**Figure 1.**
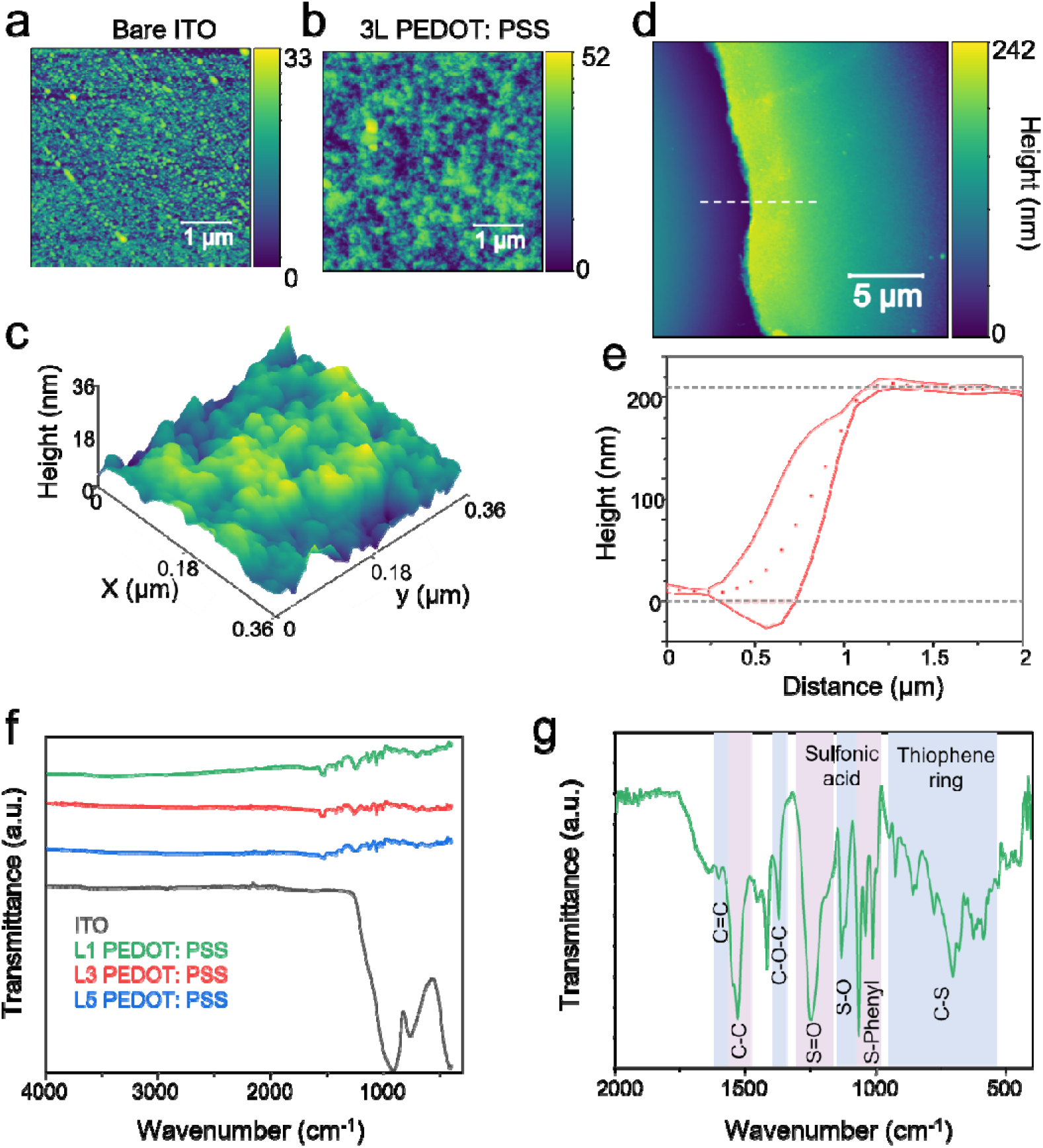
Topographical and chemical characterization of PEDOT:PSS films. (a-c) Surface topography characterized by atomic force microscopy (AFM) in air using tapping mode. Representative height retrace images of (a) bare ITO-coated glass and (b) PEDOT:PSS after three rounds of dip coating (L3 PEDOT:PSS). (c) Three-dimensional representation of a smaller scan area of the L3 PEDOT:PSS film, revealing its granular surface morphology. (d) AFM height image across a scratched region of the L3 PEDOT:PSS film used for film-thickness determination; the dashed line indicates the direction of the height-profile analysis. (e) Corresponding depth profile obtained from 10 cross-sections across the scratched region. (f) FTIR spectra of bare ITO and PEDOT:PSS-coated substrates prepared with one, three, and five dip-coating cycles (L1, L3, and L5 PEDOT:PSS, respectively). (g) Representative FTIR spectrum of PEDOT:PSS showing the characteristic vibrational bands and their assignments to the principal PEDOT and PSS chemical groups.

AFM revealed continuous PEDOT:PSS coatings with heterogeneous local thickness across the electrode surface(Figure 1a,b, Figure S1F). Films prepared with one, three, and five deposition cycles displayed comparable granular surface features despite the heterogeneous polymer distribution, consistent with the characteristic PEDOT-rich/PSS-rich heterogeneous morphology reported for PEDOT:PSS films.(16) Increasing the number of deposited layers therefore preserved the characteristic nanoscale morphology of the PEDOT:PSS films (Figure 1c, Figure S1). Height profile analysis revealed a representative film thickness of approximately 240 nm after three deposition cycles (Figure 1d,e). FTIR spectroscopy showed the characteristic vibrational bands associated with both the PEDOT backbone and the PSS dopant across all deposited layers (Figure 1f,g). The characteristic sulfonate vibrations (1260 cm^-1^, S = O; 1190 cm^-1^, S-O; S - Phenyl, 1080 cm^-1^), together with the thiophene ring vibrations, confirmed the presence of both the conducting polymer backbone and the dopant within the molecular network (Figure 1g).(17-20)

The heterogeneous morphology of the PEDOT:PSS films is particularly relevant to their potential electromechanical function. Electrochemical actuation of such a heterogeneous interface could generate localized material displacement at the cell-material interface (Scheme 1). We therefore next investigated whether electrochemical modulation of these films produces measurable deformation and whether this deformation occurs uniformly or reflects the heterogeneous polymer distribution. Although electroactive polymers such as polypyrrole have previously been incorporated into engineered microactuator devices for mechanical stimulation of cells,(21) these approaches rely on dedicated actuator architectures rather than on the intrinsic deformation of the bioelectronic interface itself. Whether electrochemical deformation of PEDOT:PSS can directly contribute to mechanosensitive signaling in cells adhered to the conducting polymer interface has remained unexplored.

### Electrochemical reduction drives structural deformation of PEDOT:PSS

We next investigated whether electrochemical modulation induces structural changes in PEDOT:PSS under conditions relevant to cell stimulation. Electrochemical characterization of PEDOT:PSS-coated electrodes was performed in a horizontal three-electrode configuration using a platinum wire as counter electrode and an Ag/AgCl reference electrode in 10 mM KCl electrolyte (Figure S2). Cyclic voltammetry recorded between -0.8 and 0.8 V showed the characteristic pseudocapacitive response of PEDOT:PSS (Figure 2a). Increasing the number of deposited layers progressively increased the enclosed voltammetric area, consistent with increased charge-storage capacity at higher polymer content. Within the cathodic potential range from approximately -0.4 to -0.8 V, a broad reduction feature was observed, consistent with reversible de-doping of PEDOT and associated structural reorganization of the PEDOT:PSS network.(5, 11) During reduction, charge compensation by solvated cations entering the polymer matrix has been associated with volumetric expansion of PEDOT:PSS.(5)

**Figure 2.**
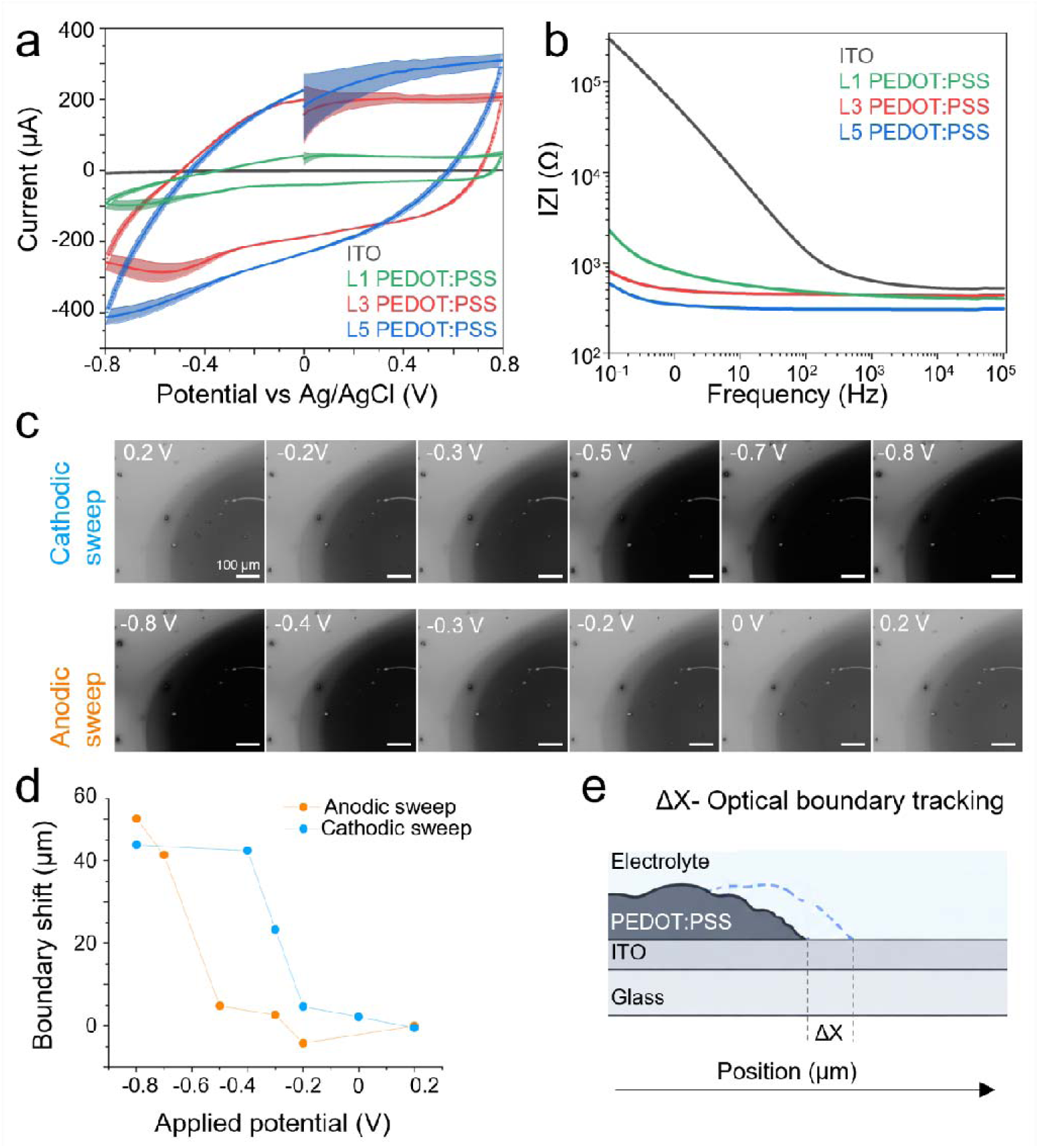
Electrochemical characterization and voltage-dependent boundary displacement of PEDOT:PSS films. (a) Cyclic voltammograms of PEDOT:PSS films comprising one (L1), three (L3), or five (L5) deposited layers, compared with bare ITO, recorded over the potential range from -0.8 to 0.8 V. Five consecutive scans were performed for each condition; solid lines represent the mean and shaded regions the standard deviation. (b) Electrochemical impedance spectra of L1, L3, and L5 PEDOT:PSS films and bare ITO over the measured frequency range. (c) Representative bright-field microscopy images of a three-layer (L3) PEDOT:PSS film acquired during cathodic and anodic potential sweeps. Images correspond to the indicated applied potentials and illustrate the voltage-dependent change in the position and optical appearance of the PEDOT:PSS/electrolyte boundary. Scale bars, 100 µm. (d) Boundary displacement extracted from the bright-field microscopy recordings as a function of applied potential during cathodic and anodic sweeps, relative to the initial boundary position. (e) Schematic representation of the boundary-tracking approach used to quantify the voltage-dependent change in the position of the PEDOT:PSS/electrolyte interface (ΔX).

Electrochemical impedance spectroscopy further showed a progressive decrease in impedance with increasing PEDOT:PSS layer number compared with bare ITO (Figure 2b), consistent with the increasing electrochemically active polymer content. As the electrochemical responses of three- and five-layer films were comparable, while three-layer coatings provided sufficient polymer coverage at lower overall thickness, three-layer PEDOT:PSS electrodes were selected for subsequent cell experiments.

To directly visualize structural changes during electrochemical cycling, cyclic voltammetry was performed simultaneously with bright-field imaging (Figure 2c and Supporting Video 1). Importantly, the observed changes were not restricted to variations in optical contrast. The position of the PEDOT:PSS/electrolyte boundary shifted with the applied potential, providing direct spatial evidence of material displacement. Tracking the boundary position revealed potential-dependent displacement during the cathodic sweep, followed by movement toward the initial position upon reversal of the potential (Figure 2d,e). These measurements demonstrate that electrochemical modulation of PEDOT:PSS produces potential-dependent microscopic deformation associated with changes in its electrochemical state.

Comparable deformation was observed for films with different numbers of deposited layers (Figure S2b,c). The response, however, was spatially non-uniform and reflected the heterogeneous distribution of the polymer. Structural changes developed more rapidly in polymer-dense regions and subsequently propagated toward less densely coated regions and the film boundary. This behavior is consistent with local redistribution of the PEDOT:PSS network during electrochemical actuation and indicates that deformation of the interface occurs locally rather than as uniform expansion of the entire film.

Temporal changes in normalized pixel intensity provided a complementary optical readout of this process (Figure S3). Within the potential interval from 0 to approximately -0.4 V, normalized pixel intensity varied approximately linearly with the applied potential, indicating that the optical response of PEDOT:PSS develops already at potentials less negative than the broad reduction feature observed by cyclic voltammetry. Together with the directly measured boundary displacement, these observations show that the material response extends into a lower-amplitude potential regime than would be inferred from the broad voltammetric reduction feature alone.

Based on these observations, -240 mV was selected as the stimulation potential for the subsequent cell experiments, as it reproducibly induced the material response while remaining within this lower-amplitude potential regime. Together, the electrochemical and optical measurements establish that PEDOT:PSS undergoes spatially heterogeneous deformation under conditions compatible with cell stimulation.

### Electrochemical actuation of PEDOT:PSS induces mechanosensitive calcium responses in HEK293T cells

Three-layer PEDOT:PSS electrodes were used as substrates for HEK293T cell cultures. Cells were grown to approximately 70% confluency and labeled with a calcium-sensitive fluorescent probe to visualize intracellular Ca^2+^ changes during stimulation (Figure 3a,b). HEK293T cells adhered directly to the PEDOT:PSS interface.

**Figure 3.**
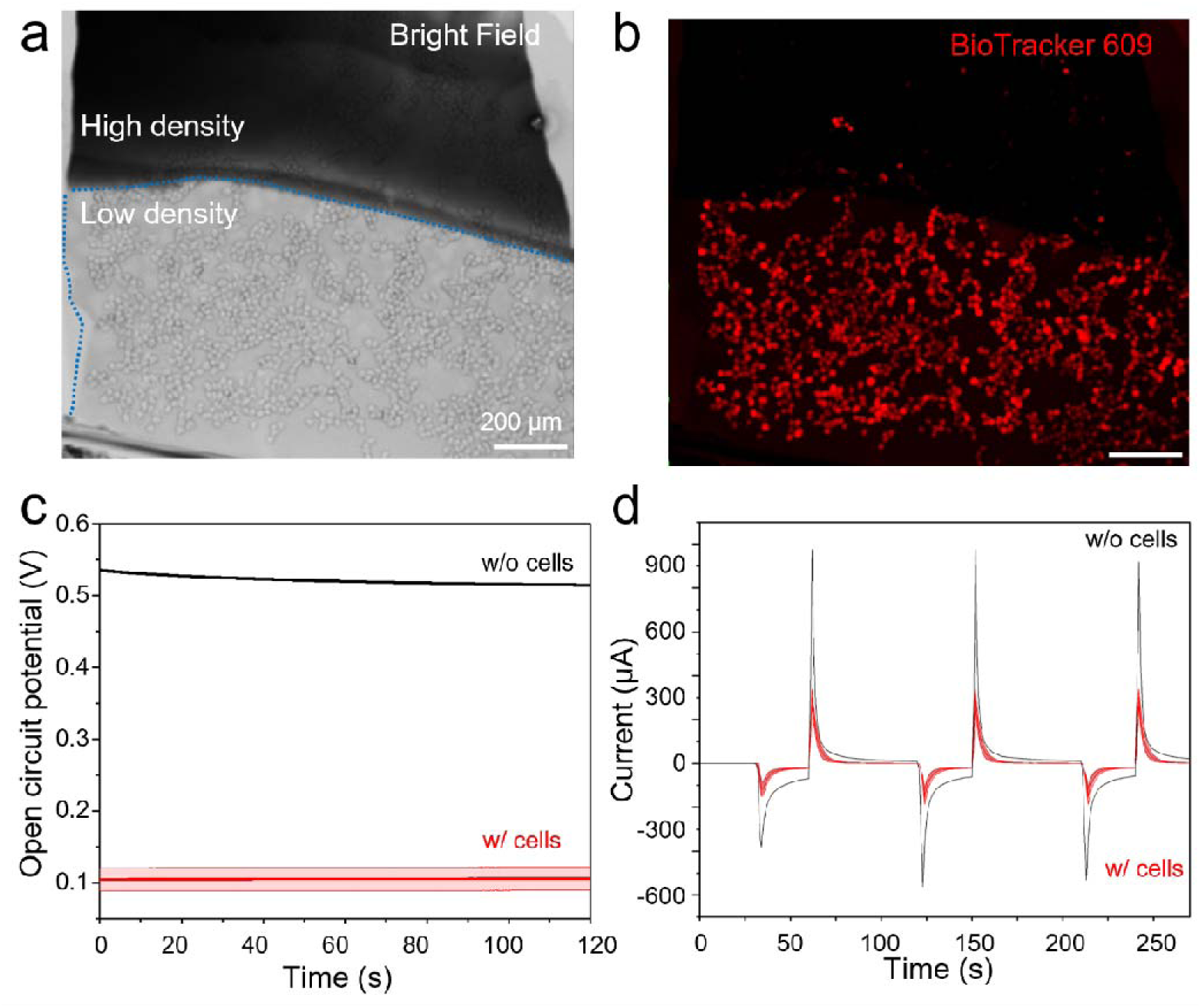
Electrical characterization of PEDOT:PSS films in the presence of HEK293T cells. (a) Representative bright-field image of HEK293T cells cultured on a transparent three-layer (L3) PEDOT:PSS film, showing regions of high and low cell density across the film surface. (b) Corresponding fluorescence image of HEK293T cells labeled with BioTracker 609 for visualization during calcium imaging experiments. Scale bars in (a) and (b), 200 µm. (c) Open-circuit potential (OCP) of L3 PEDOT:PSS electrodes measured for 120 s in Tyrode’s solution in the presence (w/ cells) and absence (w/o cells) of HEK293T cells. (d) Representative chronoamperometric current responses of L3 PEDOT:PSS electrodes with and without HEK293T cells during three consecutive stimulation cycles. The potential was stepped from the respective resting potential (∼120 mV with cells and ∼550 mV without cells) to -240 mV for 30 s per stimulation cycle.

To establish the electrochemical conditions for cell stimulation, open-circuit potentiometry (OCP) was measured in Tyrode’s solution (1.8 mM Ca^2+^) using PEDOT:PSS electrodes with and without cells (Figure 3c). The presence of cells shifted the OCP from approximately 500-550 mV for cell-free electrodes to approximately 120 mV for cell-covered electrodes, indicating a substantial change in the electrochemical equilibrium at the PEDOT:PSS/electrolyte interface. Such sensitivity of PEDOT:PSS/electrolyte interfaces to adherent HEK293T cells is consistent with previous electrochemical measurements of PEDOT:PSS-based biointerfaces.(22) The experimentally determined OCP was subsequently used as the resting potential during stimulation. Multi-step amperometry was performed by alternating 30 s at the resting potential with 30 s at -240 mV over three consecutive cycles (Figure 3d). The presence of cells altered the corresponding current response; nevertheless, reproducible electrochemical transitions were maintained throughout the three stimulation cycles. The -240 mV potential was selected as a reproducible working condition at which PEDOT:PSS deformation was observed while remaining within the lower amplitude potential regime identified in Figure 2.

The same stimulation sequence was subsequently applied during live cell calcium imaging of HEK293T cells cultured directly on PEDOT:PSS electrodes (Figure 4). Representative frames show an increase in intracellular Ca^2+^ fluorescence during application of -240 mV (Figure 4a). Analysis of 300 randomly selected cells from six independent experiments showed calcium responses across the cell population during repeated electrochemical modulation (Figure 4b). The corresponding population-averaged ΔF/F_0_ trace showed reproducible calcium transients over consecutive stimulation cycles (Figure 4c).

**Figure 4.**
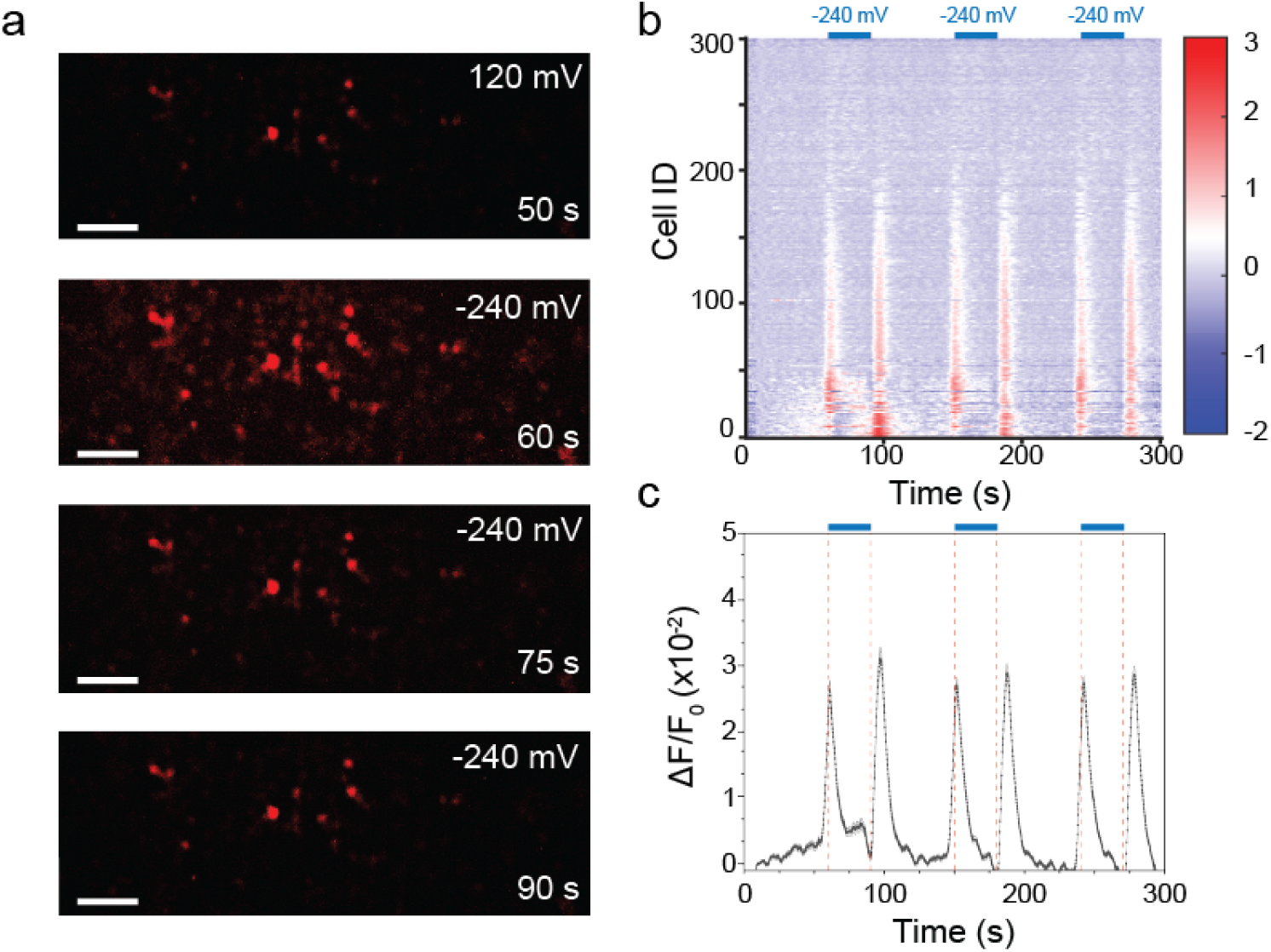
Electrochemical stimulation of PEDOT:PSS elicits calcium responses in HEK293T cells. (a) Representative frames from the live-cell calcium imaging video showing intracellular calcium fluorescence before (120 mV) and during electrochemical stimulation (-240 mV). The corresponding acquisition times are indicated in each image. Scale bars, 100 µm. (b) Heat map of ΔF/F_0_ responses from 300 randomly selected HEK293T cells. Blue bars indicate the application of a reducing potential (-240 mV). (c) Population-averaged calcium response (ΔF/F_0_) showing reproducible transient increases during each stimulation cycle.

The calcium response developed over several seconds following the electrochemical transition. Calcium transients were also observed upon restoration of the electrode to the resting potential, indicating that cellular activation was associated with transitions between electrochemical states rather than exclusively with the applied negative potential. This temporal behavior followed the deformation of PEDOT:PSS observed during electrochemical modulation (Figure 2), suggesting that the cellular response may be coupled to structural changes at the cell-polymer interface.

To test whether mechanosensitive pathways contributed to the cellular response, we employed two complementary pharmacological approaches. HEK293T cultures were incubated with GsMTx4, an inhibitor of mechanosensitive ion channels, including PIEZO1(23, 24) or with tetrodotoxin (TTX), a blocker of voltage-gated sodium channels.(25) Endogenous expression of PIEZO1 and TRPV4 in the HEK293T cultures used here was additionally confirmed by immunofluorescence (Figure S4).

GsMTx4 treatment attenuated the calcium response during electrochemical stimulation, as evident from both the individual-cell heat map and the population-averaged ΔF/F_0_ trace (Figure 5a,c). In contrast, TTX-treated cells retained pronounced calcium responses over consecutive stimulation cycles (Figure 5b,d). Direct comparison of the population-averaged traces further showed attenuation of the response in the presence of GsMTx4, whereas the response in TTX-treated cells remained closer to that of untreated HEK293T cells (Figure 5e).

**Figure 5.**
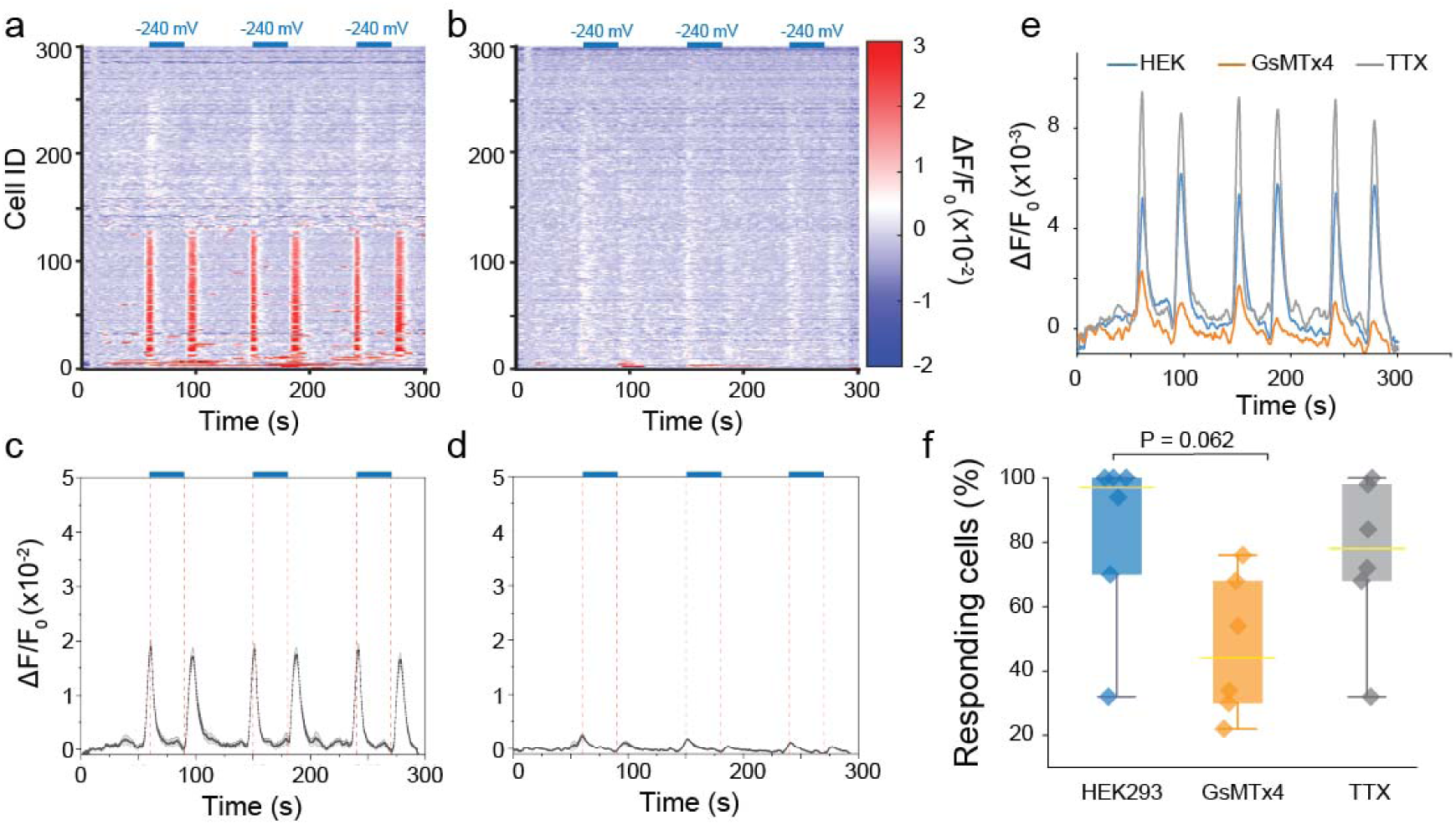
Pharmacological interrogation of electrochemically induced calcium responses supports mechanosensitive activation of HEK293T cells. (a,b) Heat maps showing ΔF/F_0_ responses of 300 randomly selected HEK293T cells following treatment with (a) 1µM GsMTx4 or (b) 1 µM tetrodotoxin (TTX). Blue bars indicate the application of -240 mV potential. (c,d) Corresponding population-averaged ΔF/F_0_ traces showing attenuation of calcium responses by GsMTx4, whereas TTX treatment largely preserves the response. (e) Overlay of the averaged ΔF/F_0_ traces from untreated HEK293T cells, GsMTx4-treated cells, and TTX-treated cells. (f) Percentage of responding cells across six independent experiments (50 randomly selected cells per experiment). GsMTx4 reduced the fraction of responding cells, whereas TTX produced a response comparable to untreated controls. The overall Kruskal-Wallis test did not reach statistical significance (P = 0.057). Pairwise comparisons using Dunn’s multiple-comparisons test with Holm correction gave P = 0.062 for HEK293T versus GsMTx4, P = 0.531 for HEK293T versus TTX, and P = 0.183 for GsMTx4 versus TTX.

To quantify this response across independent experiments, 50 cells were randomly selected from each of six experiments per condition and classified as responding when their binned ΔF/F_0_ signal exceeded five times the standard deviation of the prestimulation baseline. GsMTx4 reduced the fraction of responding cells compared with untreated HEK293T cells, although the difference did not reach statistical significance (P = 0.062), while TTX produced a responder fraction comparable to untreated cells (Figure 5f).

Together, the attenuation of the calcium response by GsMTx4, its persistence in the presence of TTX, and the temporal correspondence between cellular activation and PEDOT:PSS deformation support the involvement of mechanosensitive pathways in the observed response. Electrochemical deformation of the heterogeneous PEDOT:PSS interface can therefore provide a mechanical component to cell stimulation, in addition to its established electrical function.

### Mechanistic interpretation

Collectively, these findings support a mechanism in which electrochemical modulation of PEDOT:PSS is converted into mechanical stimulation at the cell-material interface. Electrochemical reduction drives ion and water uptake within the PEDOT:PSS network,(2, 11) accompanied here by potential-dependent microscopic deformation and displacement of the polymer film (Figure 2). Importantly, this response was spatially non-uniform, with changes occurring more rapidly in polymer-dense regions and propagating toward the film boundary. Together with the granular and heterogeneous morphology identified by AFM (Figure 1), these observations suggest that electrochemical actuation produces localized surface displacement rather than uniform expansion of the entire interface.

For cells adhered directly to the polymer, such localized displacement could transfer mechanical strain to the plasma membrane, as proposed in Scheme 1. This interpretation is supported by the calcium responses accompanying repeated electrochemical actuation (Figure 4), their attenuation following inhibition of mechanosensitive ion channels with GsMTx4, and their persistence following blockade of voltage-gated sodium channels with TTX (Figure 5). The pharmacological experiments identify the involvement of mechanosensitive pathways in the cellular response. Although the contribution of individual mechanosensitive channels was not determined in this study, immunofluorescence confirmed the endogenous expression of PIEZO1 and TRPV4 in the HEK293T cells used here (Figure S4).

Previous studies have shown that PEDOT:PSS undergoes reversible electrochemically driven swelling associated with ion exchange and changes in polymer hydration.(5, 11) Related mechanical stimulation has also been demonstrated using polypyrrole-based microactuator systems, where electrochemical deformation of an engineered actuator was used to mechanically stimulate epithelial cells(21). Here, however, mechanical stimulation arises directly from electrochemical deformation of the PEDOT:PSS biointerface itself, without the requirement for an additional actuator architecture. This establishes an electromechanical function of PEDOT:PSS in which the same material used as the conductive cell interface also provides the mechanically active component.

The ability to convert relatively small changes in electrochemical potential into local mechanical deformation expands the functional role of PEDOT:PSS beyond electrical conduction and recording. Importantly, our results show that electrochemical stimulation at a PEDOT:PSS-cell interface can include a mechanical component capable of engaging cellular mechanosensitive pathways. This contribution should be considered when interpreting cellular responses to conducting-polymer-based electrical stimulation, particularly when cells are in direct contact with the polymer interface. Control over film morphology, thickness, composition, and electrochemical response could therefore be used to tune the spatial distribution of mechanically active regions. Such interfaces could complement conventional electrical stimulation by enabling access to endogenous mechanotransduction pathways at conducting polymer–cell interfaces.

## CONCLUSIONS

In this work, we demonstrate that electrochemical actuation of PEDOT:PSS can generate mechanical cues sufficient to induce calcium responses in adherent HEK293T cells. PEDOT:PSS films exhibited a heterogeneous granular morphology and underwent potential-dependent microscopic deformation, including measurable displacement of the film boundary during electrochemical modulation. These structural changes occurred non-uniformly across the film, consistent with local redistribution of the polymer network during actuation. When HEK293T cells were cultured directly on the PEDOT:PSS interface, repeated electrochemical stimulation at -240 mV produced reproducible intracellular calcium responses. Attenuation of these responses by GsMTx4, together with their persistence in the presence of TTX, supports the involvement of mechanosensitive pathways rather than a response arising solely from voltage-gated sodium-channel excitation. The pharmacological data, together with the expression of PIEZO1 and TRPV4 in HEK293T cells, support activation of mechanosensitive pathways without assigning the response to a single molecular channel.

These findings identify PEDOT:PSS as an electromechanical biointerface in which electrochemical modulation can introduce a mechanical component alongside its established electrical function. This mechanical contribution should therefore be considered when interpreting cellular responses to conducting-polymer-based electrical stimulation. The same structural dynamics could also be exploited to couple electrochemical control with cellular mechanotransduction. Control over film morphology, composition, and electrochemical properties may allow the magnitude and spatial distribution of these mechanical cues to be tuned, providing bioelectronic interfaces in which electrical and mechanical stimulation are controlled through the same material.

## ASSOCIATED CONTENT

### Supporting Information

The Supporting Information is available free of charge.

#### Supporting Information

Additional AFM characterization of PEDOT:PSS films; electrochemical cell configuration and potential-dependent morphological changes; correlation of PEDOT:PSS deformation with cyclic voltammetry; and immunofluorescence labeling of TRPV4 and PIEZO1 in HEK293T cells.

#### Supporting Video

Bright-field microscopy video showing the dynamic morphological response of the heterogeneous PEDOT:PSS film during cyclic variation of the applied potential.

## AUTHOR INFORMATION

### Corresponding Authors

**Danijela Gregurec**, Biointerfaces lab, Department of Chemistry and Pharmacy, Friedrich-Alexander-Universitaet Erlangen-Nuernberg;

**Vicente Duran-Toro,** Biointerfaces lab, Department of Chemistry and Pharmacy, Friedrich-Alexander-Universitaet Erlangen-Nuernberg;

### Author Contributions

The manuscript was written through contributions of all authors. All authors have given approval to the final version of the manuscript.

### Notes

The Authors declare no conflict of interest.

**Scheme 1.**
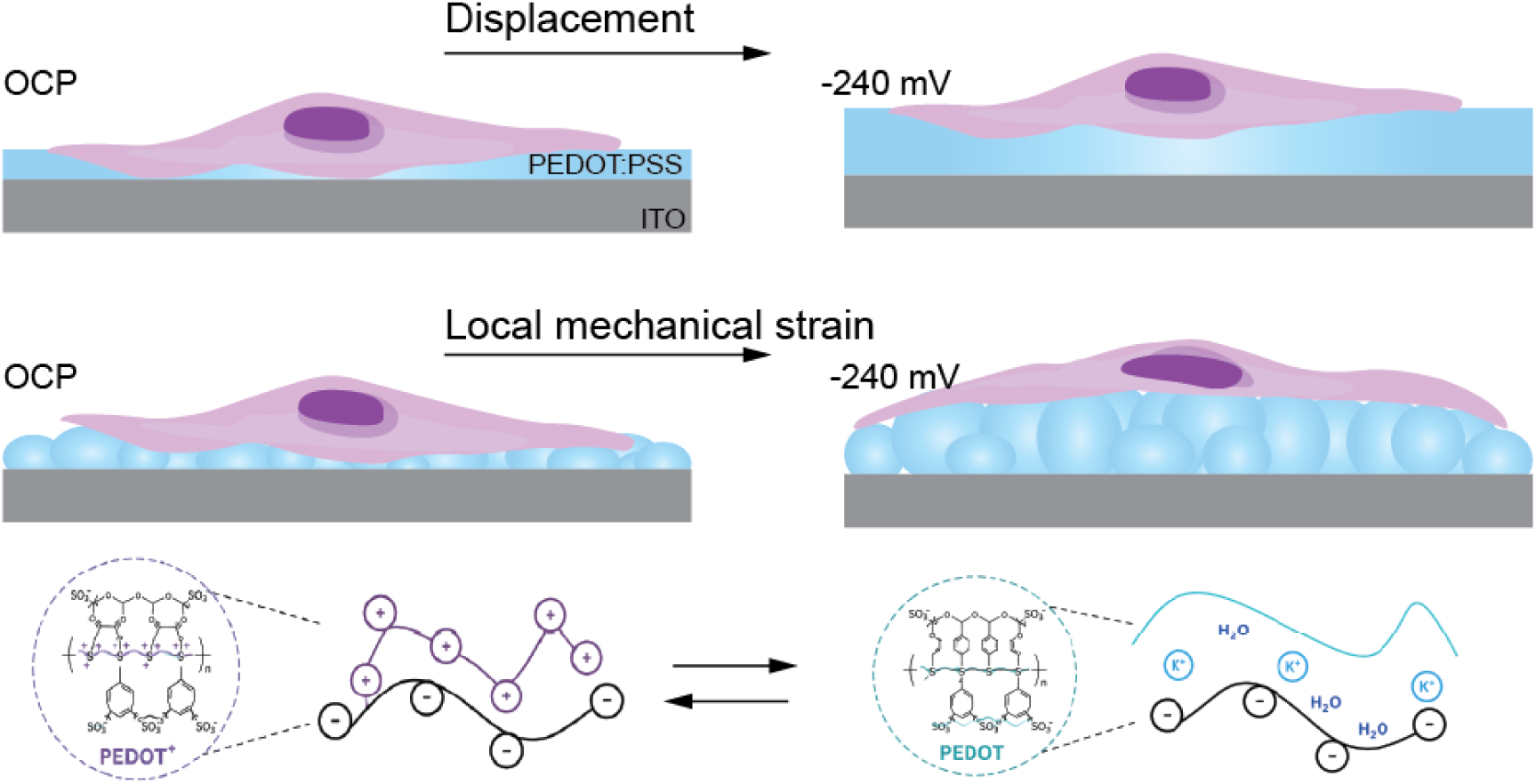
Electrochemically driven deformation of the PEDOT:PSS biointerface. Schematic representation of PEDOT:PSS deformation upon application of -240 mV relative to the open-circuit potential (OCP). Electrochemical reduction and associated ion and water redistribution induce displacement of the PEDOT:PSS film, while its heterogeneous morphology results in localized deformation at the cell-material interface that can mechanically couple to adherent cells.

## Abbreviations

AFM: atomic force microscopy
ATR: attenuated total reflection
CV: cyclic voltammetry
DBSA: dodecylbenzenesulfonic acid
DMEM: Dulbecco’s modified Eagle’s medium
EIS: electrochemical impedance spectroscopy
FBS: fetal bovine serum
FTIR: Fourier transform infrared spectroscopy
GOPS: (3-glycidyloxypropyl)trimethoxysilane
ITO: indium tin oxide
OCP: open-circuit potential
PEDOT:PSS: poly(3,4-ethylenedioxythiophene):polystyrene sulfonate
PBS: phosphate-buffered saline
ROI: region of interest
TTX: tetrodotoxin.

