## Supplementary Information for "Electrochemical Deformation of PEDOT:PSS Drives Mechanosensitive Cell Activation"

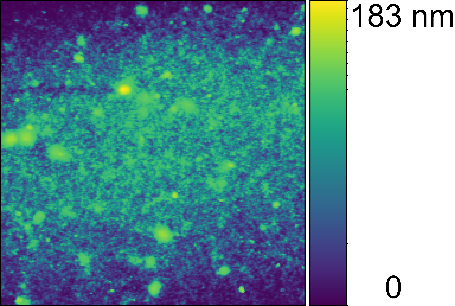

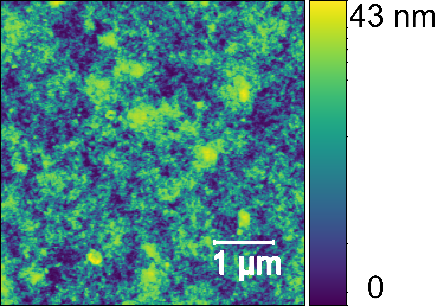

a

b

**Figure S1. Atomic force microscopy of PEDOT:PSS films deposited onto ITO glass electrodes by dip coating.** Representative AFM topography images of (a) 1-layer and (b) 5-layer PEDOT:PSS films. Measurements were performed in air using a Track60 probe (spring constant, 1.6 N/m) in tapping mode.

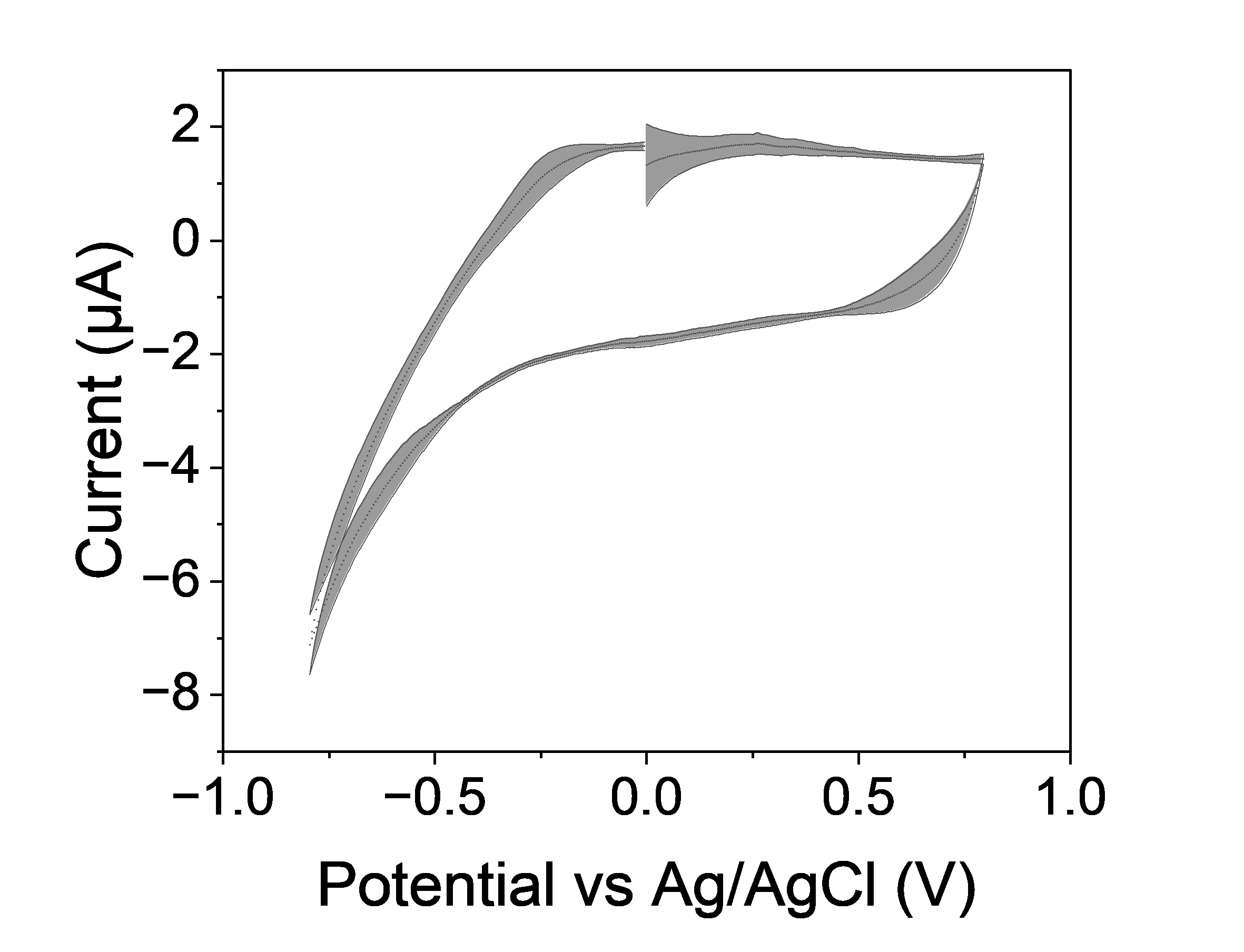

a

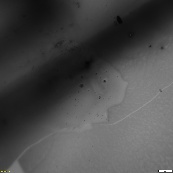

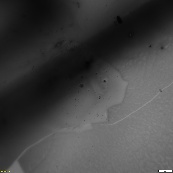

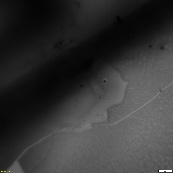

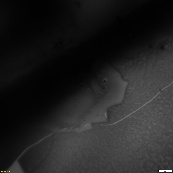

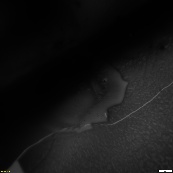

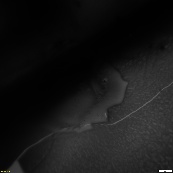

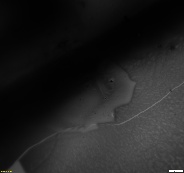

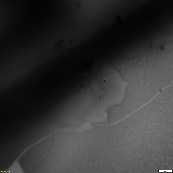

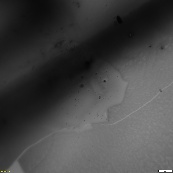

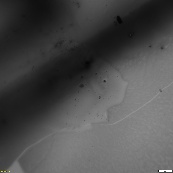

0.2 V

0.0 V

-0.2 V

-0.4 V

-0.6 V

-0.8 V

-0.6 V

-0.4 V

-0.2 V

0.0 V

c

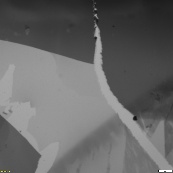

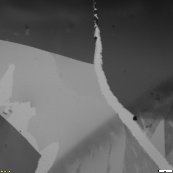

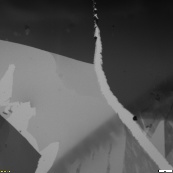

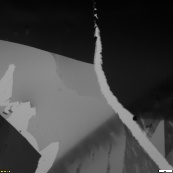

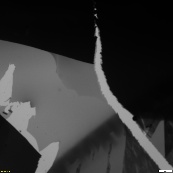

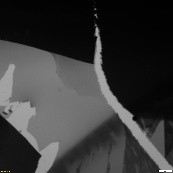

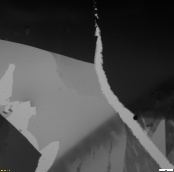

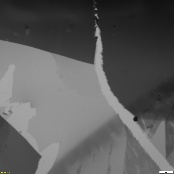

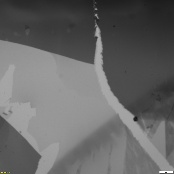

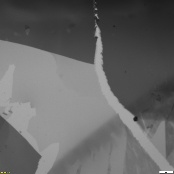

0.2 V

0.0 V

-0.2 V

-0.4 V

-0.6 V

-0.8 V

-0.6 V

-0.4 V

-0.2 V

0.0 V

b

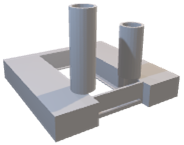

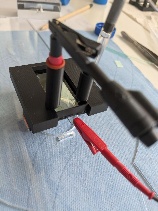

**Figure S2. Electrochemical cell configuration and voltage-dependent morphological response of PEDOT:PSS films.** (a) Rendered model of the PLA-based three-electrode electrochemical cell (left), photograph of the assembled prototype showing the horizontal slot used to position the working electrode in contact with the electrolyte (middle), and cyclic voltammogram of bare ITO glass used as the working electrode in the horizontal configuration (right). (b,c) Representative bright-field video frames acquired during cyclic voltammetry of (b) 1-layer and (c) 5-layer PEDOT:PSS films deposited onto ITO glass electrodes. The applied potentials corresponding to the individual frames are indicated.

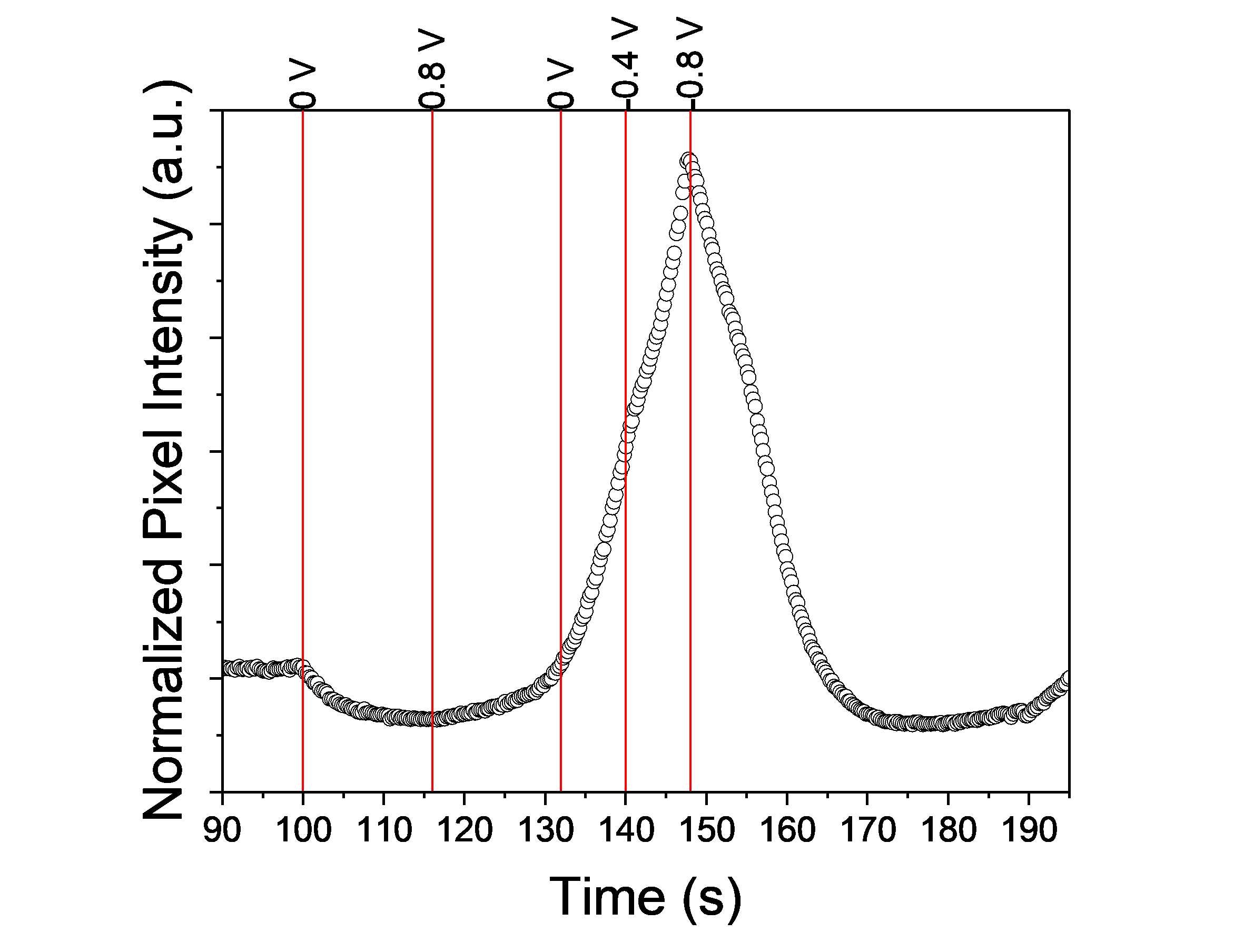
**Figure S3. Correlation between normalized pixel intensity and applied potential in a 3-layer PEDOT:PSS film.** Pixel intensity was quantified from defined regions of interest in bright-field video recordings acquired during cyclic voltammetry and normalized to background regions. Frame timestamps were aligned with the corresponding voltammogram to assign the applied potential to each time point.

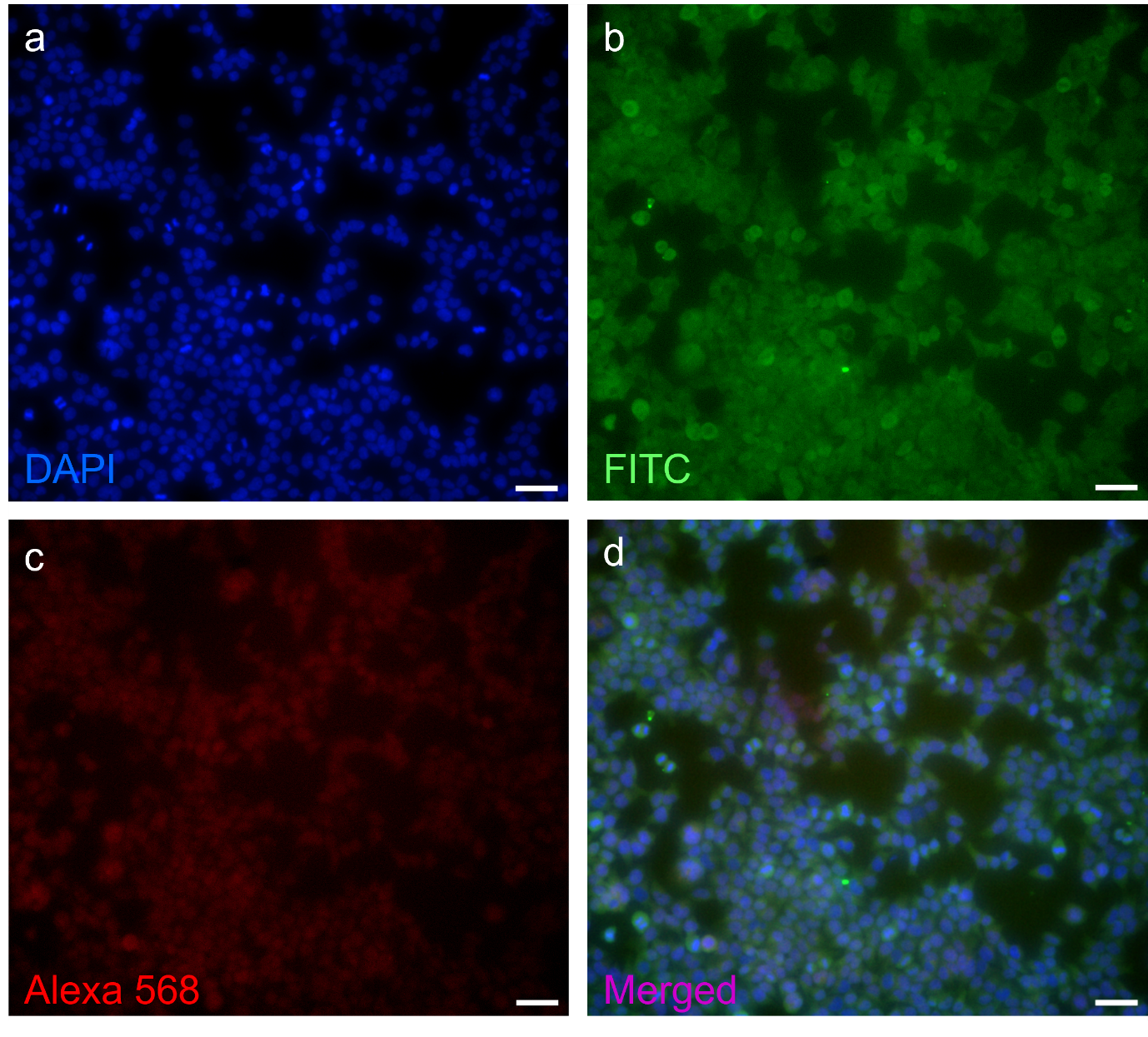

**Figure S4. Immunofluorescent labelling of TRPV4 and Piezo1 in HEK293 cells.** Representative immunofluorescence images acquired at 20× magnification showing (a) DAPI-labelled nuclei (blue), (b) TRPV4 immunolabelling detected in the green fluorescence channel, (c) Piezo1 immunolabelling detected in the red fluorescence channel, and (d) merged DAPI, TRPV4, and Piezo1 fluorescence channels. Scale bars 50 µm..

Supplementary Video

**Supplementary Video 1. Dynamic morphological response of PEDOT:PSS during cyclic variation of the applied potential.** Bright-field microscopy video of the heterogeneous PEDOT:PSS film acquired during cyclic voltammetry. Morphological changes become apparent at reducing potentials, developing preferentially in regions of higher local polymer density and propagating toward less dense regions of the film.
